# Effect of Varying Cigarette Smoke Concentrations and Durations on the Retinal Pigment Epithelium

**DOI:** 10.64898/2026.10.29.762941

**Authors:** Alyssa Chen, Wei-Hsuan Lee, Suchana Das, Inara Jawed, Ganesh Satyanarayana, Rabina Kumpakha, Sayantan Datta

## Abstract

Dry age-related macular degeneration (AMD) is a leading cause of blindness in the elderly, and treatment options are limited. Cigarette smoke (CS), the primary modifiable environmental risk factor for AMD, damages the retinal pigment epithelium (RPE) by inducing oxidative stress, which in turn activates multiple stress response pathways. Low levels of stress induce autophagy, a cell-recycling mechanism; moderate stress induces epithelial-mesenchymal transition (EMT); and high, sustained stress, once other cytoprotective pathways are exhausted, leads to cell death. All three pathways are implicated in AMD pathogenesis. Although CS is known to induce oxidative stress, its impact on these distinct stress response pathways remains understudied, and few studies have directly compared mouse and human RPE responses to the same stressor. Here, we examined the effect of varying concentrations and durations of cigarette smoke condensate (CSC) exposure on oxidative stress, autophagy, EMT, and cell death in ARPE-19 (human) and mouse RPE (MRPE) cells. Understanding which stress response pathways predominate at which CSC exposure levels may help guide the development of AMD treatments stratified by smoking history.

**NEW & NOTEWORTHY:** Cigarette smoke is the leading environmental risk factor for AMD, for which no treatment currently exists. The cellular-level impact of cigarette smoke on the RPE remains understudied. Here, we show that RPE cells demonstrate heterogeneous responses to varying levels and durations of cigarette smoke, which may advance AMD treatments personalized to individual smoking level.

## INTRODUCTION

Age-related macular degeneration (AMD) is a leading cause of irreversible blindness among the elderly with limited treatment options(1–3). Dry AMD is the most common form of the disease, accounting for approximately 85–90% of all AMD cases, and is marked by progressive loss of central vision(4) through drusen formation and degeneration of photoreceptors and the retinal pigment epithelium (RPE) (5,6), a single layer of polygonal cells located between the photoreceptor outer segments and the vascular choroid(7).

AMD is a multifactorial disease shaped by biological aging, genetic predisposition, and environmental influences, with cigarette smoke (CS) established as the leading modifiable environmental risk factor(8–10). CS has previously been linked to RPE damage and retinal disease, with a dose-dependent relationship reported between pack-years smoked and retinal disease severity(11,12).

In the RPE, CS induces oxidative stress, an imbalance between the production and elimination of reactive oxygen species (ROS)(13). Overproduction of ROS causes irreversible cell damage and contributes to AMD pathogenesis. Mitochondria are both a major source of ROS and a primary site of oxidative stress within the cell, and mitochondrial stress and dysfunction are increasingly recognized as important contributors to AMD pathophysiology. Nuclear factor erythroid 2-related factor 2 (NRF2), a key antioxidant transcription factor, regulates the antioxidant response by binding on antioxidant response elements (AREs) to induce target antioxidant genes(14,15). Aged retina shows a decreased NRF2 response to oxidative insult(16), yet little is known about how varying CS exposures affect this response. Oxidative stress also activates adenosine monophosphate-activated protein kinase (AMPK), a major regulator of metabolic homeostasis(17), converting it to its phosphorylated form (pAMPK); several studies suggest pAMPK directly activates NRF2 via phosphorylation(18,19). Increases in pAMPK to AMPK ratio indicate increased AMPK activation, which may therefore indicate an increased need to upregulate cellular responses against oxidative stress and energy imbalance. Because impaired antioxidant response in the RPE has been observed in AMD models, understanding how CS drives oxidative stress is important for clarifying its role in AMD pathogenesis.

Oxidative stress activates several distinct stress response pathways in the RPE, including autophagy, epithelial-mesenchymal transition (EMT), and cell death (20–22), each of which is dysregulated in AMD pathogenesis. These pathways likely represent an escalating hierarchy of cellular responses, in which the balance between adaptive mechanisms such as autophagy and EMT, versus outright cell death, depends on the intensity and duration of the stressor. However, the duration and level of CS exposure at which each of these distinct responses is induced remains underexplored, highlighting the need to evaluate how CS influences RPE stress response and, in turn, AMD pathogenesis.

Autophagy, a clearance mechanism in which lysosomes remove harmful cellular contents to preserve cell health, is considered one of the RPE’s least damaging responses to stress(23–26). Autophagy is carried out by several components, notably microtubule-associated protein 1A/1B-light chain 3 (LC3), which converts from its cytosolic form (LC3-I) to its membrane-associated, phosphatidylethanolamine-conjugated form (LC3-II) upon activation(27,28). Sequestosome 1 (p62), a ubiquitin-binding protein, binds ubiquitinated cargo and LC3, bridging the marking and degradation of damaged material(29). Previous studies have found that Lc3 and p62 expression varies with autophagy induction in the RPE(30,31). Autophagic regulation is delicate: optimal activation supports retinal health, while excessive activation can trigger cell death(32). Autophagic flux is impaired by cigarette smoke extract exposure in non-ocular tissue, particularly lung(33–36), and while RPE autophagy is a recognized driver of AMD, the effect of AMD risk factors such as CS on autophagic flux in the RPE remains underexplored (37).

Following prolonged oxidative stress or other moderate stressors, RPE cells can undergo epithelial-mesenchymal transition (EMT)(38,39), losing polarity and cell-cell adhesion to become mesenchymal cells(40,41). This process, commonly seen in cancer metastasis, pulmonary fibrosis, and subretinal fibrosis in AMD and other degenerative retinal diseases(41–43), produces the elongated, spindle-shaped morphology characteristic of migratory mesenchymal cells(44). EMT is marked by increased expression of the transcription factors SNAIL1 and ZEB1(45), increased expression of the mesenchymal marker vimentin, and decreased expression of the epithelial marker E-cadherin (46). As a transcriptional program, EMT allows cells to survive stress at the cost of their specialized epithelial function(41); in the RPE, EMT contributes to the development of subretinal fibrosis, a hallmark of late-stage AMD, though little is known about how CS-induced oxidative stress influences this process. RPE cells with mitochondrial dysfunction undergo EMT through retrograde mitochondria-to-nucleus signaling(47), and mice acutely exposed to CSC show enrichment of differentially expressed genes related to EMT and senescence, along with an increased proportion of dedifferentiated RPE cells whose transcriptome reflects a mix of up- and downregulated genes involved in mitochondrial function and metabolism(48).

When RPE cells face acute, high-level stress that exhausts other cytoprotective mechanisms, they undergo cell death. Death of RPE cells is a primary characteristic of late-stage AMD known as geographic atrophy(49). Although CS is known to cause cell death, the effect of varying CS durations and dosages on RPE cell death has not been characterized(50).

While previous studies have examined the effect of CS on other RPE processes, including differentiation and immune response, in both mouse and human models(48,51,52), no study to date has investigated how varying levels and durations of CS affect stress response pathways in the RPE, nor directly compared mouse and human models, a comparison that is critical for guiding model selection and translation to humans. To address this gap, we evaluated the effect of CSC on RPE stress response pathways, including oxidative stress, autophagy, EMT, and eventual cell death, across a range of doses (0, 100, 250, and 500 µg/ml) and two timepoints (4 and 24 hours) in both ARPE-19 (human) and mouse RPE (MRPE) cells.

## MATERIALS AND METHODS

### Cell Culture

ARPE-19 (ATCC, VA, USA) and immortalized mouse RPE (MRPE) (Applied Biological Materials, BC, Canada) cells were maintained in DMEM-F12 with 10% fetal bovine serum (FBS) and in PriGrow III with 10% fetal bovine serum (FBS) media, respectively. Cells were seeded at 50,000 cells/cm2 and grown to 70-80% confluency before experimentation. For RNA and protein extraction, cells were plated in 12-well plates; for imaging studies, cells were plated in 96-well plates or in 8-chamber slides. Cells were maintained at 37°C in a humidified atmosphere of 5% CO2, with media exchanged twice weekly.

### CSC Treatment

40 mg/ml stock CSC (Murty Pharmaceuticals) was diluted in serum-free media to generate three treatment concentrations: 100, 250, and 500 μg/ml. DMSO at the same final concentration as CSC 250 μg/ml was used as the vehicle control. Cells were treated at each concentration in biological triplicate at 80% confluency for 4 or 24 hours.

### RT-qPCR

Total RNA was extracted using the Quick-RNA Miniprep Kit (Zymo Research), quantified on Take3 (Biotek) and 100 ng of RNA was reverse transcribed using the High-Capacity RNA to cDNA Kit (Thermo Fisher Scientific). 0.5 ul of cDNA were used per PCR reaction (10 µl), performed on a CFX Opus Real-Time PCR System (Bio-Rad) using TaqMan (Thermo Fisher Scientific) primer probes (Supplemental Table 1) under the manufacturer’s default thermal cycling conditions (40 cycles of 1 s at 95°C and 20 s at 60°C). GAPDH served as the reference gene, and results were expressed as fold change relative to control using the ddCt method.

### Western Blot

ARPE-19 and MRPE cells were washed twice with PBS following each treatment. Protein extracts were prepared using RIPA lysis buffer (ThermoFisher) with protease and phosphatase inhibitor cocktail, and protein concentration was measured using a DC protein assay (Bio-Rad). Samples (5 or 10 µg protein) were prepared in NuPAGE LDS sample buffer (4X, Invitrogen), separated by SDS-PAGE using 4–12% NuPAGE Bis-Tris gels (ThermoFisher), and transferred to nitrocellulose membranes using the Bio-Rad Trans-Blot Turbo Transfer System. Membranes were blocked for one hour in 5% BSA or 5% non-fat milk in Tris-buffered saline with 0.1% Tween-20 (TBST), incubated overnight at 4°C with primary antibody (Supplemental Table 2), and then incubated with the appropriate HRP-conjugated secondary antibody (1:2000), with three TBST washes between steps. Signals were detected using a chemiluminescence detection system (Pierce ECL or SuperSignal West Femto, Thermo Fisher Scientific) and imaged on a ChemiDoc MP Imaging System (Bio-Rad). Band intensities are reported as arbitrary densitometric units.

### Measurement of Viability and Cell Morphology

Cell death was visualized using ethidium homodimer-1 (EthD-1, Invitrogen). Cells were washed with PBS, stained with 2 µM EthD-1 for 30 minutes at 37°C, washed again, and counterstained with 1 µM Hoechst (Invitrogen) for 5 minutes to visualize nuclei. Imaging was performed on an EVOS M5000 microscope (excitation/emission 495/635 nm for EthD-1). The number of EthD-1-positive cells was counted and expressed as a percentage of control. Brightfield imaging on the same microscope was used to assess cell morphology. Cell aspect ratio was calculated using ImageJ to evaluate morphometric changes consistent with cellular EMT.

### Mitochondrial ROS Assessment

Mitochondrial ROS was visualized using MitoSox Superoxide Red (500 nM, Life Technologies), with cells counterstained with 1 µM Hoechst (Invitrogen) to visualize nuclei. Cells were incubated with MitoSox for 30 minutes at 37°C, and images were acquired on an EVOS M5000 microscope (excitation/emission 396/610 nm). Mitochondrial ROS was quantified by fluorescence intensity and normalized to control and to cell number.

### Gene and Protein Nomenclature

Targets are referred to in the text by their common names (for example NRF2, p62, LC3 and SNAIL1). The official human and mouse gene symbols and the corresponding protein names are given in Supplemental Table 3.

### Statistics and Sample Size

All experiments were performed in three biological replicates with three technical replicates where applicable. Following confirmation of normal distribution, two-tailed, heteroscedastic Student’s t-tests were used to establish statistical significance at α = 0.05.

## RESULTS

### CSC increases mitochondrial superoxide levels in ARPE-19 and MRPE at both 4 and 24 hours in a concentration-dependent manner

As mitochondria are a major site of cellular ROS production, we examined the effect of varying CSC concentrations and durations on mitochondrial superoxide levels. In ARPE-19 cells, superoxide levels increased with CSC concentration at both 4 and 24 hours of treatment (Figures 1A, B); at 4 hours, superoxide levels were significantly higher at CSC 500 compared to control (p < 0.001). In MRPE cells, superoxide levels also increased in a concentration-dependent manner at both timepoints, with all treatment groups differing significantly from control (Figures 1C, D).

**Figure 1.**
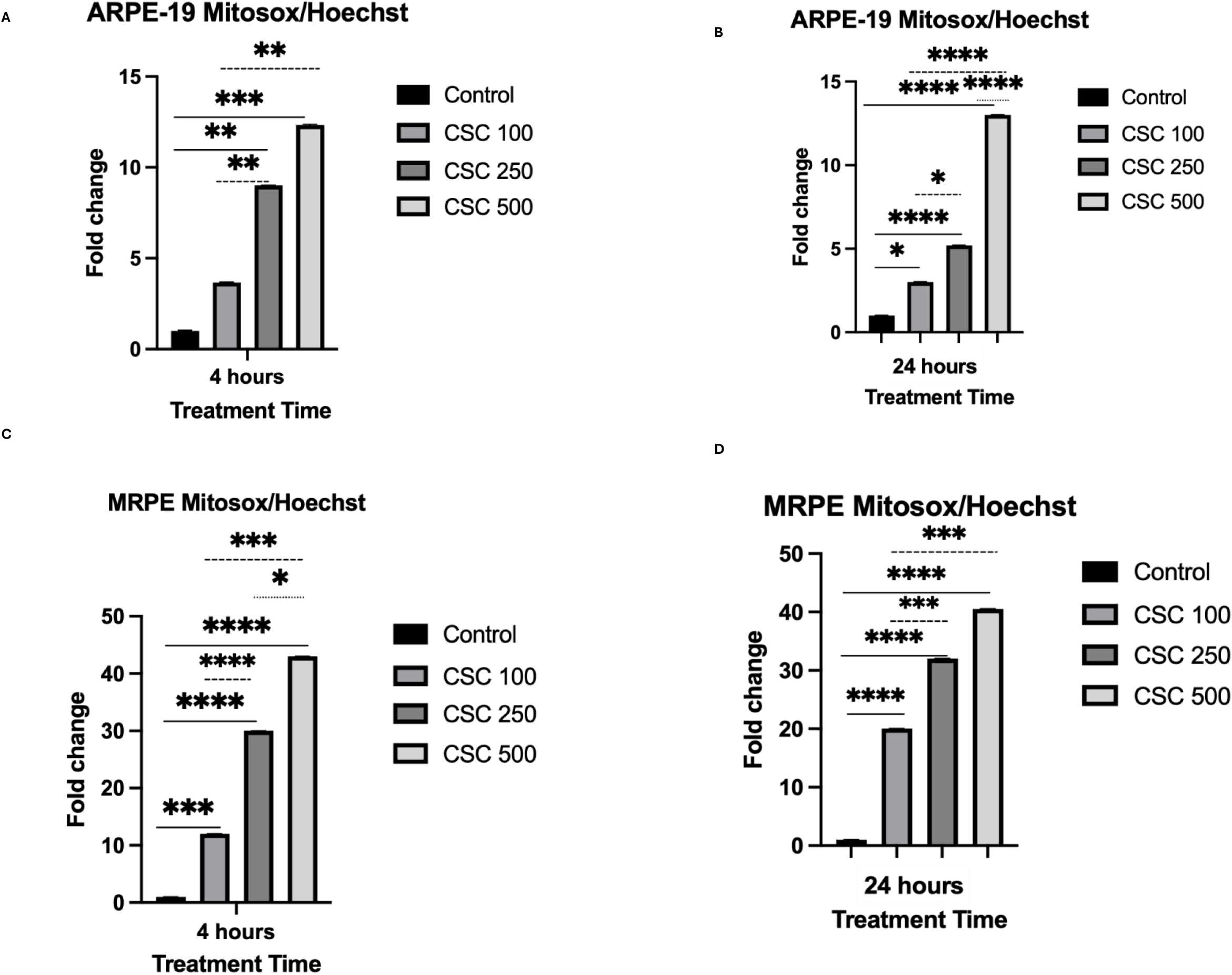
CSC increases mitochondrial superoxide levels in a concentration-dependent manner in ARPE-19 (A, B) and MRPE (C, D) cells at 4 and 24 hours of CSC treatment. Mitosox (red) and Hoechst (blue) stain in ARPE-19 and mouse RPE cells treated for 4 and 24 hours with increasing concentrations of CSC (0, 100, 250, 500 ug/mL) imaged with EVOS microscope at 10X magnification. DMSO is used as control. The number of red fluorescent spots (Mitosox red) normalized to cell number (Hoechst blue) is quantified and presented in the graph (N=6 per treatment condition). Fold change compared to the control was calculated at each concentration for 3 biological replicates per concentration. Statistical comparisons were performed using Student’s t-test. * p<0.05, ** p<0.01, *** p<0.001, ****p < 0.0001.

Though trends were similar between the two cell types, magnitude differed: at both timepoints, superoxide levels increased to nearly 15-fold in ARPE-19 cells but over 40-fold in MRPE cells, suggesting the two cell types differ in sensitivity to CSC. These trends were consistent with EVOS imaging (Supplemental Figure 1).

### CSC induces concentration-dependent oxidative stress response in both ARPE-19 and MRPE at 4 and 24 hours of CSC exposure

Having established that CSC increases mitochondrial superoxide in a concentration-dependent manner, we next examined its effect on oxidative stress response pathways.

At 4 hours, qPCR analysis in ARPE-19 showed a significant increase in NRF2 expression between control and CSC 100 (p = 0.007) and between control and CSC 250 (p = 0.033), followed by a decrease between CSC 250 and CSC 500 (p = 0.177) (Figure 2A). MRPE showed a similar pattern, with a significant increase in Nrf2 expression at CSC 250 (p = 0.019) followed by a decrease at CSC 500 (Figure 2B). These results suggest an initial concentration-dependent upregulation of NRF2 up to CSC 250, with a diminished response at higher CSC concentrations.

**Figure 2.**
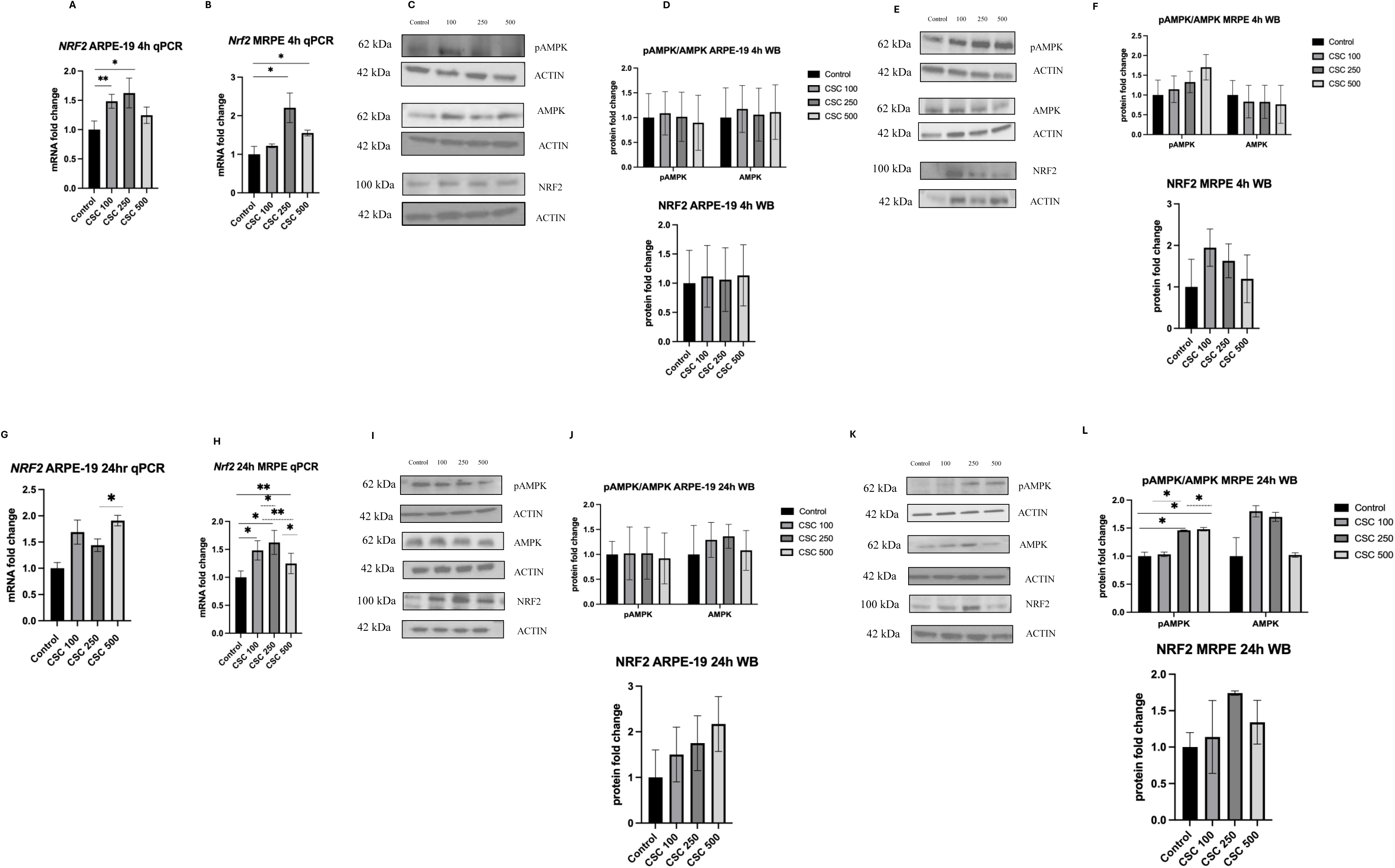
4 and 24 hours of CSC treatment induces oxidative stress response in ARPE-19 and mouse RPE. NRF2 mRNA expression quantified using TaqMan RT-qPCR and plotted as fold change after ddCT calculations (N = 3) in A) ARPE-19 treated for 4 hours and B) MRPE treated for 4 hours. Western blot analysis of pAMPK, AMPK, Nrf2, and beta-actin in C) ARPE-19 treated for 4 hours with D) corresponding quantification, and E) MRPE treated for 4 hours with F) corresponding quantification. NRF2 mRNA expression quantified using TaqMan RT-qPCR and plotted as fold change after ddCT calculations (N = 3) in G) ARPE-19 treated for 24 hours and H) MRPE treated for 24 hours. Western blot analysis of pAMPK, AMPK, Nrf2, and beta-actin in I) ARPE-19 treated for 24 hours with J) corresponding quantification, and in K) MRPE treated for 24 hours with L) corresponding quantification. Band intensities quantified with Image J, normalized to beta-actin, and plotted as fold change. Fold change is represented in mean ± SD, Student’s t-test; * p<0.05, ** p<0.01, *** p<0.001, ****p < 0.0001.

Western blot analysis of pAMPK and AMPK in ARPE-19 at 4 hours showed little change across treatment groups, with a small increase in pAMPK between control and CSC 100 followed by decreases at higher concentrations; AMPK and NRF2 protein levels were similarly unchanged across groups (Figures 2C, D). In MRPE at 4 hours, pAMPK protein increased and AMPK protein decreased with increasing CSC concentration (Figures 2E, F), consistent with a concentration-dependent oxidative stress response. Additionally, NRF2 protein followed trends similar to that of NRF2 qPCR analysis (Figures 2E, F).

At 24 hours, qPCR analysis in ARPE-19 showed an increase in NRF2 expression between control and CSC 100, a decrease between CSC 100 and CSC 250, and a significant increase between CSC 250 and CSC 500 (p = 0.041) (Figure 2G); overall, NRF2 expression rose with CSC concentration. In MRPE, Nrf2 expression followed a pattern similar to ARPE-19 at 4 hours, increasing between control and CSC 250 (p = 0.001) before decreasing significantly between CSC 250 and CSC 500 (p = 0.011) (Figure 2H).

Western blot analysis in ARPE-19 at 24 hours showed unchanged pAMPK across groups, while AMPK increased before decreasing between CSC 250 and CSC 500; NRF2 protein increased with CSC concentration, mirroring the corresponding qPCR trend (Figures 2I, J). In MRPE at 24 hours, pAMPK protein increased between CSC 100 and CSC 250, AMPK protein increased between control and CSC 100 before decreasing at higher concentrations, and NRF2 protein showed an initial increase followed by a decrease, similar to the trend in ARPE-19 (Figures 2K, L).

### CSC induces concentration-dependent autophagy stress responses at transcriptional levels in ARPE-19 and MRPE at 4 hours and 24 hours of CSC treatment

Because autophagy is an early-stage response to stressors such as oxidative stress, we examined the effect of CSC on the autophagy markers LC3 and P62.

At 4 hours, qPCR analysis in ARPE-19 showed a significant, concentration-dependent increase in LC3 expression, while P62 increased between control and CSC 250 before decreasing between CSC 250 and CSC 500 (Figure 3A). MRPE showed a similar pattern: a concentration-dependent increase in Lc3, and an increase in p62 between control and CSC 250 followed by a decrease at CSC 500 (Figure 3B). In both cell types, LC3/Lc3 was significantly upregulated between control and CSC 500 (ARPE-19, p = 0.0002; MRPE, p = 0.0096), and P62/p62 was significantly downregulated between CSC 250 and CSC 500 (ARPE-19, p = 0.0276; MRPE, p = 0.0359).

**Figure 3.**
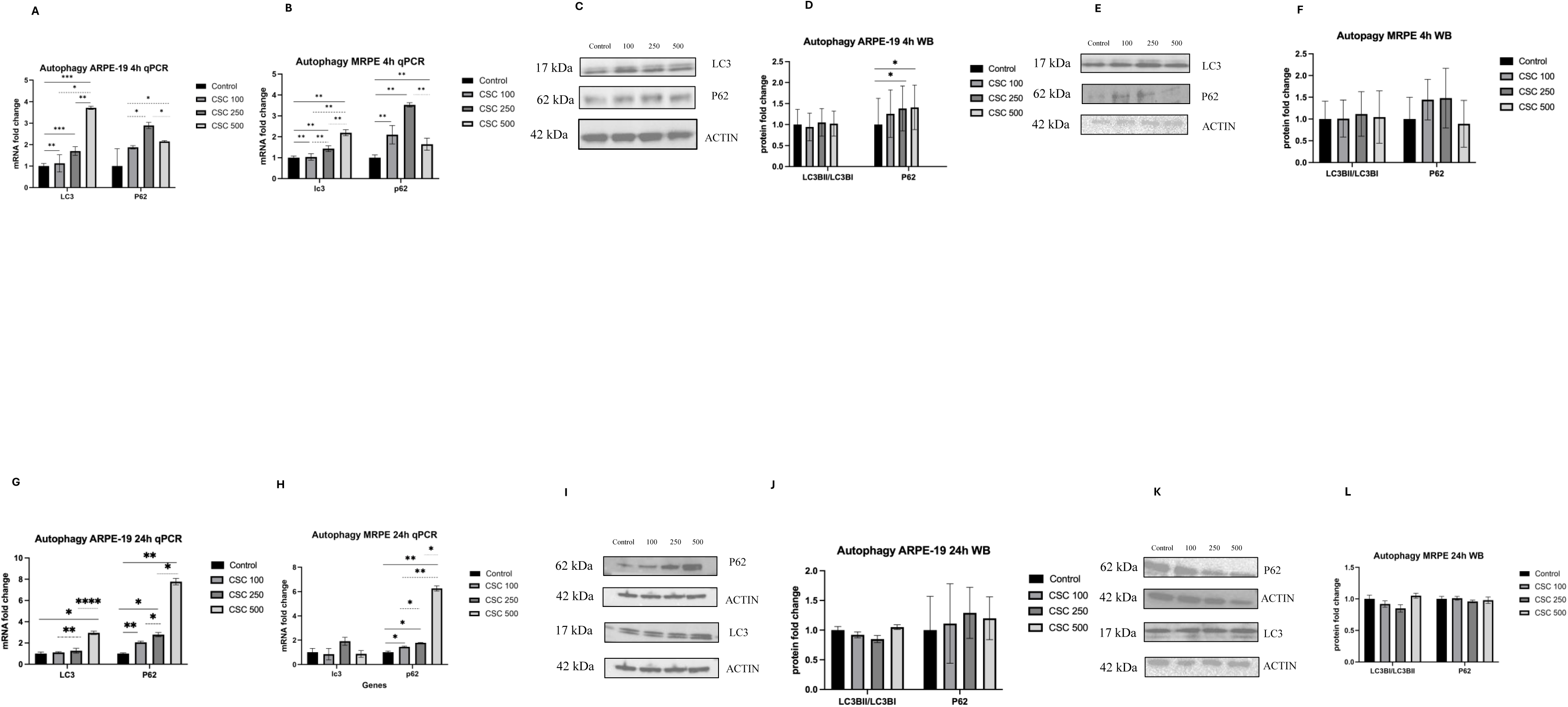
CSC changes expression of autophagy related genes at 4 hours and 24 hours in both ARPE-19 and MRPE. LC3 and P62 mRNA expression quantified using TaqMan RT-qPCR and plotted as fold change after ddCT calculations (N = 3) in A) ARPE-19 treated for 4 hours and B) MRPE treated for 4 hours. Western blot analysis of LC3, P62, and beta-actin in C) ARPE-19 treated for 4 hours with D) corresponding quantification, and E) MRPE treated for 4 hours with F) corresponding quantification. LC3 and P62 mRNA expression quantified using TaqMan RT-qPCR and plotted as fold change after ddCT calculations (N = 3) in G) ARPE-19 treated for 24 hours and H) MRPE treated for 24 hours. Western blot analysis of LC3, P62, and beta-actin in I) ARPE-19 treated for 24 hours with J) corresponding quantification, and in K) MRPE treated for 24 hours with L) corresponding quantification. Band intensities quantified with Image J, normalized to beta-actin, and plotted as fold change. 5 ug protein was loaded in each sample. C) F) I) L): Band intensities quantified with Image J, normalized to beta-actin, and plotted as fold change. Fold change is represented in mean ± SD, Student’s t-test; * p<0.05, ** p<0.01, *** p<0.001, ****p < 0.0001.

Western blot analysis of LC3B-I-to-LC3B-II turnover in ARPE-19 at 4 hours showed no consistent trend across treatment groups, while P62 protein increased with CSC concentration (Figures 3C, D). In MRPE at 4 hours, LC3B turnover showed no significant change, while P62 protein increased with CSC concentration before decreasing between CSC 250 and CSC 500 (Figures 3E, F).

At 24 hours, qPCR analysis in ARPE-19 showed a dose-dependent increase in LC3, consistent with the 4-hour trend; unlike at 4 hours, P62 increased with CSC concentration throughout (Figure 3G). In MRPE, p62 also increased in a concentration-dependent manner, while Lc3 showed no consistent directional trend (Figure 3H).

Western blot analysis in ARPE-19 at 24 hours showed largely unchanged LC3B-II/LC3B-I turnover, with a non-significant decrease between control and CSC 250 followed by an increase at CSC 500; P62 protein was similarly unchanged, with a small non-significant increase to CSC 250 followed by a decrease at CSC 500 (Figure 3I, J). MRPE showed a comparable pattern for LC3B turnover, while P62 protein remained unchanged across all groups (Figure 3K, L).

Across both cell types and timepoints, LC3 trends differed between qPCR and western blot analysis, likely because western blot captures LC3B-I-to-LC3B-II turnover specifically, while qPCR reflects total LC3 transcript levels, which may not track with this turnover. The differing P62 qPCR and western blot trends may reflect a similar distinction between transcriptional and post-translational regulation.

### CSC induces concentration-dependent EMT stress responses at transcriptional levels in ARPE-19 and MRPE at 4 hours and 24 hours of CSC treatment

EMT is a more severe response by the epithelial cells where cells undergo adaptive changes to survive sub-lethal severe stress. We examined the effect of CSC on the EMT transcription factors SNAIL1 and ZEB1, and on the EMT markers E-cadherin and Vimentin, and captured brightfield imaging to further assess EMT.

At 4 hours, qPCR analysis showed similar trends in both cell types: SNAIL1/Snai1 and ZEB1/Zeb1 both increased with CSC concentration in ARPE-19 (Figure 4A) and MRPE (Figure 4B). Western blot analysis of E-CADHERIN in ARPE-19 at 4 hours showed little change with increasing CSC concentration, while VIMENTIN increased from control to CSC 100 and from CSC 250 to CSC 500 (Figures 4C, D). In MRPE at 4 hours, E-CADHERIN showed a small, non-significant increase with CSC concentration, as did VIMENTIN bewteen CSC 100 to CSC 500 (Figures 4E, F).

**Figure 4.**
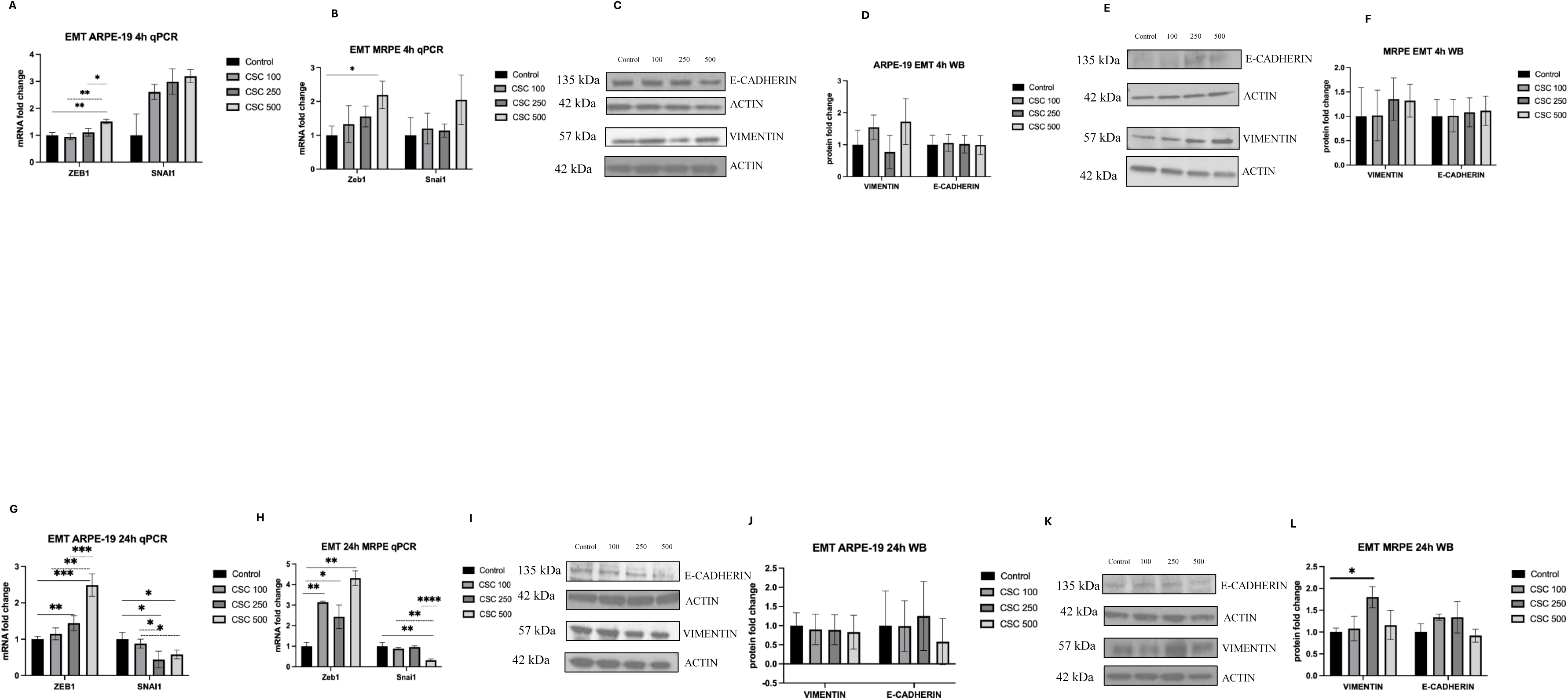
CSC induces EMT stress responses in ARPE-19 and MRPE at 24 hours transcriptionally. mRNA expression of ZEB1 and SNAIL1 were quantified using TaqMan RT-qPCR and plotted as fold change after ddCT calculations (N = 3) in A) ARPE-19 treated for 4 hours and B) MRPE treated for 4 hours. Western blot analysis of Vimentin, E-cadherin, and beta-actin in C) ARPE-19 treated for 4 hours with D) corresponding quantification, and E) MRPE treated for 4 hours with F) corresponding quantification. ZEB1 and SNAIL1 mRNA expression quantified using TaqMan RT-qPCR and plotted as fold change after ddCT calculations (N = 3) in G) ARPE-19 treated for 24 hours and H) MRPE treated for 24 hours. Western blot analysis of Vimentin, E-cadherin, and beta-actin in I) ARPE-19 treated for 24 hours with J) corresponding quantification, and in K) MRPE treated for 24 hours with L) corresponding quantification. Band intensities quantified with Image J, normalized to beta-actin, and plotted as fold change. Fold change is represented in mean ± SD, Student’s t-test; * p<0.05, ** p<0.01, *** p<0.001, ****p < 0.0001.

At 24 hours, SNAIL1/Snai1 expression followed a different pattern than at 4 hours in both cell types, trending downward overall. In ARPE-19, SNAIL1 decreased with increasing CSC concentration before rising again between CSC 250 and CSC 500 (Figure 4G); in MRPE, Snail1 decreased between control and CSC 100 and between CSC 250 and CSC 500 (Figure 4H). For both cell types, only the decrease in SNAIL1 expression between control and CSC 500 was statistically significant (p = 0.01 for ARPE-19, p = 0.008 for MRPE). ZEB1/Zeb1 expression at 24 hours followed a pattern similar to 4 hours in both cell types, increasing overall with a significant increase between control and CSC 500; in MRPE, Zeb1 showed a small, non-significant decrease between CSC 100 and CSC 250 before resuming an upward trend. In ARPE-19, VIMENTIN showed a non-significant decrease as CSC concentration increased, while E-CADHERIN showed a non-significant increase between CSC 100 and CSC 250 and a decrease between CSC 250 and CSC 500 (Figures 4I, J). Western blot analysis of MRPE at the same timepoint for vimentin showed a significant increase between control and CSC 250, and for E-CADHERIN, yielded a same trend as ARPE-19 at the same point, with a non-significant decrease between CSC 250 and CSC 500 (Figures 4K, L)

Because SNAIL1 marks early-stage EMT and ZEB1 marks late-stage EMT, these results suggest that CSC induces both early- and late-stage EMT at 4 hours in both cell types, while late-stage EMT predominates by 24 hours.

Morphometric analyses of brightfield images revealed significant elongation of cells after 24 hours of CSC treatment especially in ARPE-19 cells but also in MRPE cells but only at CSC 500 dose (Supplemental Figures 2 and 3).

Taken together, these results show that CSC induces EMT-associated transcriptional changes in both ARPE-19 and MRPE cells at 4 hours, but the effects become more pronounced after 24 hours of treatment.

### CSC leads to increased cell death in ARPE-19 and MRPE at CSC treatment for 4 hours and 24 hours in a treatment-dependent manner

Because cell death occurs once other stress responses have been exhausted, we used ethidium homodimer cell viability assays to examine the effect of CSC on cell death. In ARPE-19, cell death increased significantly with CSC concentration at both timepoints, and was higher at 24 hours than at 4 hours across at CSC 500 (Figure 5A, B). MRPE showed a similar general concentration-dependent increase in cell death at both timepoints (Figure 5C, D). These trends were consistent with EVOS imaging (Supplemental Figure 4).

**Figure 5.**
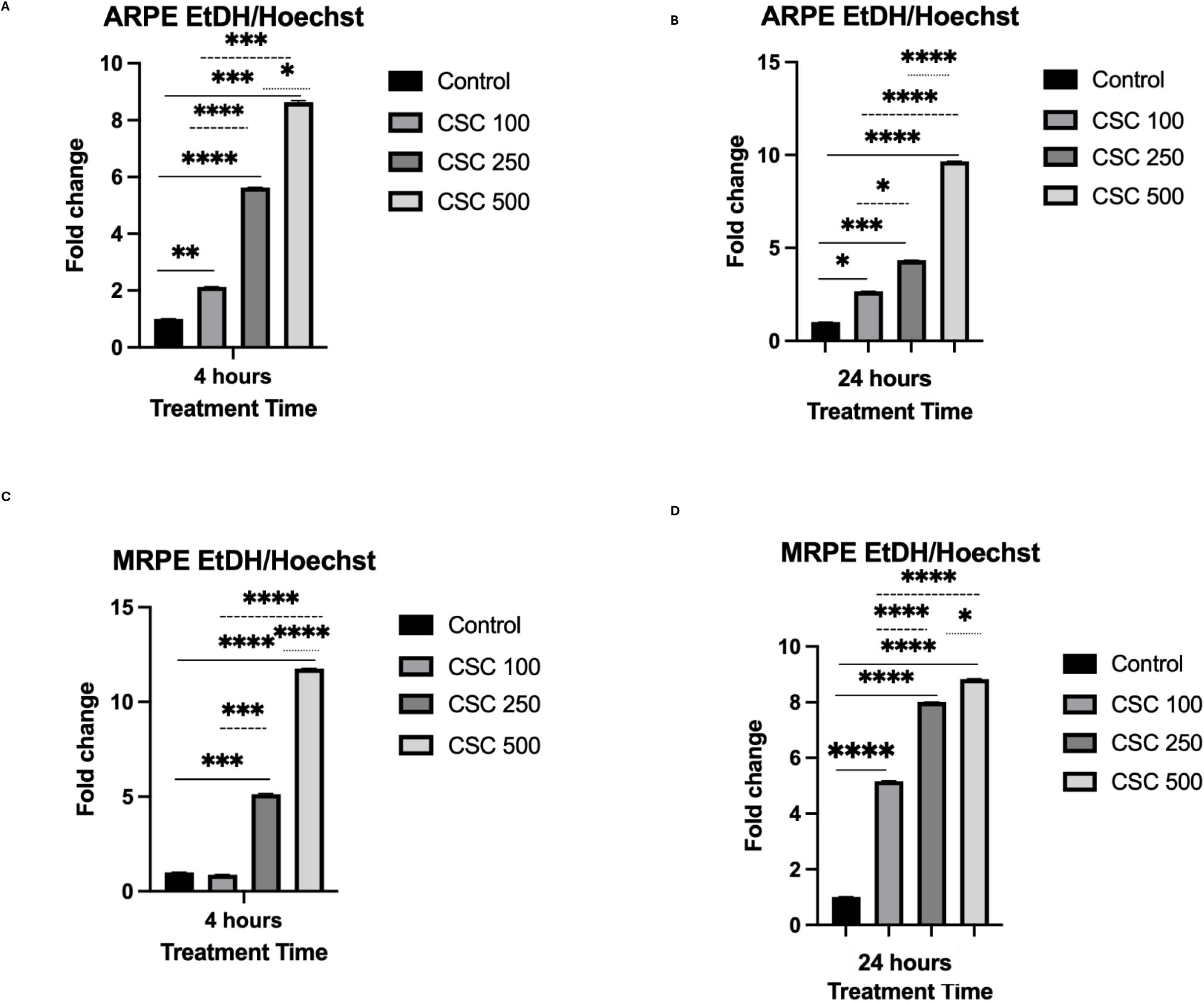
CSC increases cell death in a concentration-dependent manner in ARPE-19 and MRPE cells at 4 and 24 hours of CSC treatment. Ethidium homodimer (red) stains dead cell nuclei and Hoechst (blue) which stains live cell nuclei in ARPE-19 and mouse RPE cells treated with increasing doses of CSC at 4 hours and 24 hours. Red fluorescence was quantified and normalized to Hoechst stain. Fold change compared to the control was calculated at each concentration for 3 biological replicates per concentration (N=6 per treatment condition). Fold change is represented in mean ± SD, Student’s t-test; * p<0.05, ** p<0.01, *** p<0.001, ****p < 0.0001.

## DISCUSSION

In this study, we found that RPE cells exhibit heterogeneous responses to varying levels and durations of external oxidative stress. Autophagic responses predominated at low CSC concentrations and short exposure (100 µg/ml, 4 hours), while other cytoprotective mechanisms and cell death pathways were co-induced as CSC concentration and exposure time increased.

Within the last decade, growing evidence has shown that RPE heterogeneity, defined as the co-existence of dedifferentiated, EMT, and dying RPE cells within the same tissue, is a defining feature of AMD(47,53–56). While CS has been shown to induce RPE heterogeneity(48), the duration and level of exposure required to produce these distinct features remains underexplored, highlighting the need to understand the dose- and time-dependent effects of CSC on RPE heterogeneity and AMD risk.

To our knowledge, this is the first study to examine CSC-induced RPE heterogeneity across both varying dosages and timepoints, using the same experimental design in two cell types. Our findings show that CSC induces RPE heterogeneity in a time- and dose-dependent manner: mitochondrial superoxide assays and antioxidant response markers indicate that CSC induces oxidative stress in a treatment-dependent manner, and CSC exposure also produced treatment-dependent increases in autophagy, EMT, and cell death. The similarity of these responses across ARPE-19 and MRPE cells suggests that human and mouse RPE respond comparably to CSC.

Several patterns emerged across the stress response pathways examined in this study. For oxidative stress, CSC increased mitochondrial superoxide in both cell types at both durations, and qPCR analysis of NRF2/Nrf2 in both cell types at 4 hours showed an increase between control and CSC 250 followed by a decrease at CSC 500, indicating an oxidative stress response that peaks at intermediate CSC concentrations and subsides at the highest concentration tested. In MRPE, this pattern was mirrored in western blot analysis of NRF2 at 24 hours. Additionally, in MRPE, western blot pAMPK and AMPK trends at both durations were consistent with an oxidative stress response, with pAMPK increasing and AMPK decreasing as CSC concentration increased, though at 24 hours, AMPK decreased only after increasing between control and CSC 100(18,57). By contrast, ARPE-19 pAMPK and AMPK protein levels remained largely unchanged at both durations, suggesting a cell-line difference in this arm of the oxidative stress response.

For autophagy, both cell types showed a concentration-dependent increase in LC3/Lc3 by qPCR at both durations outside of MRPE at 24 hours, though this trend was not consistent in western blot analysis, likely reflecting the difference between total LC3 transcript levels and LC3B-I-to-LC3B-II protein turnover. At 4 hours, P62/p62 initially increased and then decreased with CSC concentration in both cell types by qPCR, while at 24 hours P62/p62 increased continuously with concentration; this shift may reflect duration-dependent differences in the autophagy response, with autophagy potentially declining at higher CSC concentrations during short exposures but continuing to rise during prolonged exposure. Together, these findings support a concentration- and duration-dependent link between CSC and autophagy, consistent with the elevated, dysregulated autophagy previously reported in AMD(58).

For EMT, both cell types showed a concentration-dependent increase in EMT markers across both durations. ZEB1/Zeb1 increased with CSC concentration by qPCR in both cell types and durations; SNAIL1/Snai1 showed a similar increase at 4 hours but not at 24 hours. Because SNAIL1 marks early-stage EMT and ZEB1 marks late-stage EMT, these results suggest that CSC induces both phases of EMT at 4 hours, while late-stage EMT predominates by 24 hours. E-cadherin and vimentin protein levels were largely unchanged by western blot at both durations, though vimentin trended upward at 24 hours in both cell types, consistent with the expected increase in vimentin during EMT. These findings indicate that CSC induces EMT in both ARPE-19 and MRPE cells at both short and prolonged exposures.

Both cell types showed a concentration-dependent increase in cell-death at both durations, based on EVOS imaging and quantification of ethidium homodimer-positive cells. Taken together, these findings show that higher CSC concentrations, at both short and prolonged exposures, are associated with upregulated oxidative stress, autophagy, EMT, and cell death. The co-occurrence of these upregulated stress response pathways alongside cell death reflects RPE heterogeneity, consistent with genetic models of AMD RPE. These results clarify the contexts in which EMT is induced in RPE and point to potentially attractive targets for treatments aimed at modifying stress-induced RPE damage.

Concentration-dependent increases in autophagy and EMT markers were evident by qPCR but not by western blot, suggesting that changes at the protein level lag transcription and may emerge only after the time points examined here.

This study has several limitations. First, we selected 4 and 24 hours to represent acute and chronic responses, but smoking behavior spans a much wider range of acuity and chronicity, and no established method exists for translating human smoking exposure into an equivalent in vitro CSC dose. Future studies should examine additional CSC concentrations and more closely spaced timepoints over a longer duration.

Second, it is difficult to estimate how well direct CSC exposure in this in vitro model reflects CS inhalation in humans. Future work using in vivo animal models in whole-body exposure chambers(46), which expose mice to CS by inhalation, could help model how CS reaches and affects the RPE systemically.

By demonstrating CSC-induced RPE heterogeneity, our findings suggest a need for AMD treatments that acknowledges multiple RPE stress responses, an approach not yet reflected in clinical practice. By showing that CSC alters expression of specific biomarkers of these processes (e.g., LC3, SNAIL1, NRF2), our findings also support ongoing research into the therapeutic potential of these biomarkers (52,59).

While the impact of smoking on the RPE remains understudied, CS remains the leading environmental risk factor for AMD(8), and understanding its impact is critical for identifying the pathogenic causes of AMD and potential therapeutic targets. Our study shows that CSC induces RPE heterogeneity, a hallmark of AMD, in a dose- and time-dependent manner, with co-induction of different responses to the same stressor in the same population of RPE. This heterogeneity reflects the complex, varied presentation of RPE in response to environmental insults such as CS. Understanding the dose- and time-dependent effects of CS may help identify which cytoprotective mechanisms dominate at which exposure levels. Future work should investigate whether pharmacological or genetic modulation of the dose-dependent thresholds identified here can prevent the transition from adaptive cytoprotection to irreversible RPE damage and EMT in preclinical models of dry AMD.

## DATA AVAILABILITY

Data generated during and/or analyzed in the manuscript are available from the corresponding author on reasonable request.

## ACKNOWLEDGMENTS

Many thanks to the NEI/NIH and the Research to Prevent Blindness Foundation for their generous funding, and to Emory University for their provision of resources critical to this work. Additional thanks to Micah Chrenek for his support and assistance.

## AUTHOR CONTRIBUTIONS

AC conceptualized the project, conducted experiments, analyzed data, made figures, and wrote and edited the manuscript. WL conducted experiments, analyzed data, made figures, and wrote and edited the manuscript. SD conducted experiments, analyzed data, and edited the manuscript. IJ helped with cigarette smoke extract preparation. GS contributed to cell culture. RK contributed to cell imaging. SD led overall idea generation and helped with manuscript writing and editing.

---

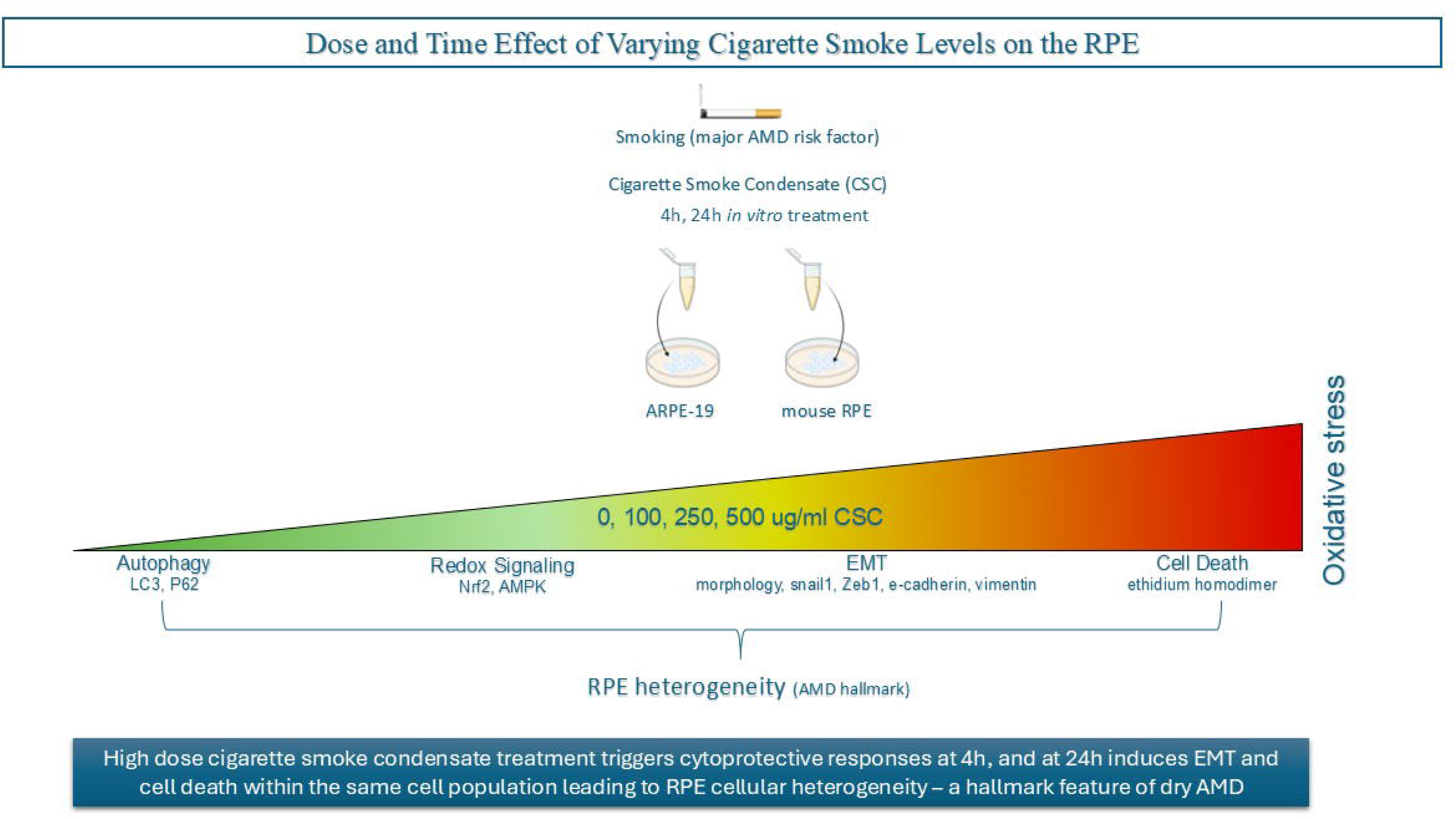

## REFERENCES

1. Gardner A, Wang S, Daniels A, Li D, Wu C, Lin L, Hong C, Zhao SR, Born RT, Kruczek K, Weist G, Barrio Real L, Wicks J, Nayal M, Carter A, Ott C, Haurigot V, Cepko CL. AAV NRF2 gene therapy preserves retinal structure and function in rodent models of oxidative damage. Mol Ther. 2026 Feb 6;S1525-0016(26)00091-2. doi:10.1016/j.ymthe.2026.02.005 PubMed PMID: 41655020.

2. Li Y, Yuan S, Zhang C, Hu P, Huang X, Zhou J, Liu D, Zhou X. Engineered nanoparticles for subconjunctival delivery to the retinal pigment epithelium: A multi-target therapy for dry AMD. J Control Release. 2026 Feb 10;390:114563. doi:10.1016/j.jconrel.2025.114563 PubMed PMID: 41435998.

3. Ye L, Zhao T, Tian H, Li R, Li Z, Li H, Liao R, Chuan J, Xu H, Zou L, Qu C, Shi Y, Yang Z, Huang L. Unveiling the choroidal immune landscape revealed interferon-gamma and TNF-alpha as novel therapeutic targets in dry AMD. Sci China Life Sci. 2026 Feb;69(2):523–41. doi:10.1007/s11427-025-2951-5 PubMed PMID: 41077603.

4. Yorston D. Age-related macular degeneration (AMD): an introduction. Community Eye Health. 2025 Jan 31;37(125):7. PubMed PMID: 40151374.

5. Al Gwairi O, Thach L, Zheng W, Osman N, Little PJ. Cellular and Molecular Pathology of Age-Related Macular Degeneration: Potential Role for Proteoglycans. J Ophthalmol. 2016;2016:2913612. doi:10.1155/2016/2913612 PubMed PMID: 27563459; PubMed Central PMCID: PMC4983667.

6. Bowes Rickman C, Farsiu S, Toth CA, Klingeborn M. Dry Age-Related Macular Degeneration: Mechanisms, Therapeutic Targets, and Imaging. Invest Ophthalmol Vis Sci. 2013 Dec;54(14):ORSF68–80. doi:10.1167/iovs.13-12757 PubMed PMID: 24335072; PubMed Central PMCID: PMC3864379.

7. Yang S, Zhou J, Li D. Functions and Diseases of the Retinal Pigment Epithelium. Front Pharmacol. 2021 Jul 28;12:727870. doi:10.3389/fphar.2021.727870 PubMed PMID: 34393803; PubMed Central PMCID: PMC8355697.

8. Cano M, Thimmalappula R, Fujihara M, Nagai N, Sporn M, Wang AL, Neufeld AH, Biswal S, Handa JT. Cigarette smoking, oxidative stress, the anti-oxidant response through Nrf2 signaling, and Age-related Macular Degeneration. Vision Research. 2010 Mar;50(7):652–64. doi:10.1016/j.visres.2009.08.018

9. Henning Y, Willbrand K, Larafa S, Weißenberg G, Matschke V, Theiss C, Görtz GE, Matschke J. Cigarette smoke causes a bioenergetic crisis in RPE cells involving the downregulation of HIF-1α under normoxia. Cell Death Discov. 2023 Oct 25;9(1):398. doi:10.1038/s41420-023-01695-5

10. Ratnapriya R, Chew EY. Age-related macular degeneration – clinical review and genetics update. Clin Genet. 2013 Aug;84(2):160–6. doi:10.1111/cge.12206 PubMed PMID: 23713713; PubMed Central PMCID: PMC3732788.

11. Khan JC, Thurlby DA, Shahid H, Clayton DG, Yates JRW, Bradley M, Moore AT, Bird AC; Genetic Factors in AMD Study. Smoking and age related macular degeneration: the number of pack years of cigarette smoking is a major determinant of risk for both geographic atrophy and choroidal neovascularisation. Br J Ophthalmol. 2006 Jan;90(1):75–80. doi:10.1136/bjo.2005.073643 PubMed PMID: 16361672; PubMed Central PMCID: PMC1856879.

12. Velilla S, García-Medina JJ, García-Layana A, Dolz-Marco R, Pons-Vázquez S, Pinazo-Durán MD, Gómez-Ulla F, Arévalo JF, Díaz-Llopis M, Gallego-Pinazo R. Smoking and age-related macular degeneration: review and update. J Ophthalmol. 2013;2013:895147. doi:10.1155/2013/895147 PubMed PMID: 24368940; PubMed Central PMCID: PMC3866712.

13. Pizzino G, Irrera N, Cucinotta M, Pallio G, Mannino F, Arcoraci V, Squadrito F, Altavilla D, Bitto A. Oxidative Stress: Harms and Benefits for Human Health. Oxid Med Cell Longev. 2017;2017:8416763. doi:10.1155/2017/8416763 PubMed PMID: 28819546; PubMed Central PMCID: PMC5551541.

14. He F, Ru X, Wen T. NRF2, a Transcription Factor for Stress Response and Beyond. Int J Mol Sci. 2020 Jul 6;21(13):4777. doi:10.3390/ijms21134777 PubMed PMID: 32640524; PubMed Central PMCID: PMC7369905.

15. Ngo V, Duennwald ML. Nrf2 and Oxidative Stress: A General Overview of Mechanisms and Implications in Human Disease. Antioxidants (Basel). 2022 Nov 27;11(12):2345. doi:10.3390/antiox11122345 PubMed PMID: 36552553; PubMed Central PMCID: PMC9774434.

16. Sachdeva MM, Cano M, Handa JT. Nrf2 signaling is Impaired in the Aging RPE given an Oxidative Insult. Exp Eye Res. 2014 Feb;119:111–4. doi:10.1016/j.exer.2013.10.024 PubMed PMID: 24216314; PubMed Central PMCID: PMC3946784.

17. Garcia D, Shaw RJ. AMPK: mechanisms of cellular energy sensing and restoration of metabolic balance. Mol Cell. 2017 Jun 15;66(6):789–800. doi:10.1016/j.molcel.2017.05.032 PubMed PMID: 28622524; PubMed Central PMCID: PMC5553560.

18. Xu W, Zhao T, Xiao H. The Implication of Oxidative Stress and AMPK-Nrf2 Antioxidative Signaling in Pneumonia Pathogenesis. Front Endocrinol (Lausanne). 2020;11:400. doi:10.3389/fendo.2020.00400 PubMed PMID: 32625169; PubMed Central PMCID: PMC7311749.

19. Joo MS, Kim WD, Lee KY, Kim JH, Koo JH, Kim SG. AMPK Facilitates Nuclear Accumulation of Nrf2 by Phosphorylating at Serine 550. Mol Cell Biol. 2016 Jul 15;36(14):1931–42. doi:10.1128/MCB.00118-16 PubMed PMID: 27161318; PubMed Central PMCID: PMC4936058.

20. Cano M, Wang L, Wan J, Barnett BP, Ebrahimi K, Qian J, Handa JT. Oxidative Stress Induces Mitochondrial Dysfunction and a Protective Unfolded Protein Response in RPE cells. Free Radic Biol Med. 2014 Apr;69:1–14. doi:10.1016/j.freeradbiomed.2014.01.004 PubMed PMID: 24434119; PubMed Central PMCID: PMC3960355.

21. Datta S, Cano M, Ebrahimi K, Wang L, Handa JT. The Impact of Oxidative Stress and Inflammation on RPE Degeneration in Non-neovascular AMD. Prog Retin Eye Res. 2017 Sep;60:201–18. doi:10.1016/j.preteyeres.2017.03.002 PubMed PMID: 28336424; PubMed Central PMCID: PMC5600827.

22. Liu J, Copland DA, Theodoropoulou S, Chiu HAA, Barba MD, Mak KW, Mack M, Nicholson LB, Dick AD. Impairing autophagy in retinal pigment epithelium leads to inflammasome activation and enhanced macrophage-mediated angiogenesis. Sci Rep. 2016 Feb 5;6(1):20639. doi:10.1038/srep20639

23. Kaarniranta K, Tokarz P, Koskela A, Paterno J, Blasiak J. Autophagy regulates death of retinal pigment epithelium cells in age-related macular degeneration. Cell Biol Toxicol. 2017;33(2):113–28. doi:10.1007/s10565-016-9371-8 PubMed PMID: 27900566; PubMed Central PMCID: PMC5325845.

24. Kocherlakota S, Das Y, Swinkels D, Vanmunster M, Callens M, Vinckier S, Vaz FM, Sinha D, Van Veldhoven PP, Fransen M, Baes M. The murine retinal pigment epithelium requires peroxisomal β-oxidation to maintain lysosomal function and prevent dedifferentiation. Proc Natl Acad Sci U S A. 120(43):e2301733120. doi:10.1073/pnas.2301733120 PubMed PMID: 37862382; PubMed Central PMCID: PMC10614831.

25. Sinha D, Valapala M, Shang P, Hose S, Grebe R, Lutty GA, Zigler JS Jr, Kaarniranta K, Handa JT. Lysosomes: regulators of autophagy in the retinal pigmented epithelium. Exp Eye Res. 2016 Mar;144:46–53. doi:10.1016/j.exer.2015.08.018 PubMed PMID: 26321509; PubMed Central PMCID: PMC4698066.

26. Wu J, Chen X, Liu X, Huang S, He C, Chen B, Liu Y. Autophagy regulates TGF-β2-induced epithelial-mesenchymal transition in human retinal pigment epithelium cells. Mol Med Rep. 2018 Mar;17(3):3607–14. doi:10.3892/mmr.2017.8360 PubMed PMID: 29286127; PubMed Central PMCID: PMC5802162.

27. Tanida I, Ueno T, Kominami E. LC3 and Autophagy. Methods Mol Biol. 2008;445:77–88. doi:10.1007/978-1-59745-157-4_4 PubMed PMID: 18425443.

28. Wang AL, Lukas TJ, Yuan M, Du N, Handa JT, Neufeld AH. Changes in Retinal Pigment Epithelium Related to Cigarette Smoke: Possible Relevance to Smoking as a Risk Factor for Age-Related Macular Degeneration. PLOS ONE. 2009 Apr 24;4(4):e5304. doi:10.1371/journal.pone.0005304

29. Komatsu M, Kageyama S, Ichimura Y. p62/SQSTM1/A170: physiology and pathology. Pharmacol Res. 2012 Dec;66(6):457–62. doi:10.1016/j.phrs.2012.07.004 PubMed PMID: 22841931.

30. Feng H, Zhao X, Guo Q, Feng Y, Ma M, Guo W, Dong X, Deng C, Li C, Song X, Han S, Cao L. Autophagy resists EMT process to maintain retinal pigment epithelium homeostasis. Int J Biol Sci. 2019 Jan 1;15(3):507–21. doi:10.7150/ijbs.30575 PubMed PMID: 30745838; PubMed Central PMCID: PMC6367589.

31. Golestaneh N, Chu Y, Xiao YY, Stoleru GL, Theos AC. Dysfunctional autophagy in RPE, a contributing factor in age-related macular degeneration. Cell Death Dis. 2018 Jan;8(1):e2537–e2537. doi:10.1038/cddis.2016.453

32. Denton D, Kumar S. Autophagy-dependent cell death. Cell Death Differ. 2019 Apr;26(4):605–16. doi:10.1038/s41418-018-0252-y PubMed PMID: 30568239; PubMed Central PMCID: PMC6460387.

33. Kim MJ, Hong J, Chang HW, Lee YH, Myong JP, Jung AR, Lee YS, Kwon M. Autophagy activation in response to cigarette smoke: Exploring the disparity in laryngeal cancer incidence and outcomes between sexes in South Korea. Transl Oncol. 2024 Dec 4;52:102229. doi:10.1016/j.tranon.2024.102229 PubMed PMID: 39637635; PubMed Central PMCID: PMC11664157.

34. Kono Y, Colley T, To M, Papaioannou AI, Mercado N, Baker JR, To Y, Abe S, Haruki K, Ito K, Barnes PJ. Cigarette smoke-induced impairment of autophagy in macrophages increases galectin-8 and inflammation. Sci Rep. 2021 Jan 11;11:335. doi:10.1038/s41598-020-79848-0 PubMed PMID: 33432024; PubMed Central PMCID: PMC7801483.

35. Vij N, Chandramani-Shivalingappa P, Van Westphal C, Hole R, Bodas M. Cigarette smoke-induced autophagy impairment accelerates lung aging, COPD-emphysema exacerbations and pathogenesis. Am J Physiol Cell Physiol. 2018 Jan 1;314(1):C73–87. doi:10.1152/ajpcell.00110.2016 PubMed PMID: 27413169; PubMed Central PMCID: PMC5866380.

36. Xu L, Li X, Wang H, Xie F, Liu H, Xie J. Cigarette smoke triggers inflammation mediated by autophagy in BEAS-2B cells. Ecotoxicol Environ Saf. 2019 Nov 30;184:109617. doi:10.1016/j.ecoenv.2019.109617 PubMed PMID: 31476449.

37. Govindaraju VK, Bodas M, Vij N. Cigarette smoke induced autophagy-impairment regulates AMD pathogenesis mechanisms in ARPE-19 cells. PLoS One. 2017 Aug 2;12(8):e0182420. doi:10.1371/journal.pone.0182420 PubMed PMID: 28767736; PubMed Central PMCID: PMC5540403.

38. Yang IH, Lee JJ, Wu PC, Kuo HK, Kuo YH, Huang HM. Oxidative stress enhanced the transforming growth factor-β2-induced epithelial-mesenchymal transition through chemokine ligand 1 on ARPE-19 cell. Sci Rep. 2020 Mar 4;10:4000. doi:10.1038/s41598-020-60785-x PubMed PMID: 32132577; PubMed Central PMCID: PMC7055234.

39. Yang YC, Chien Y, Yarmishyn AA, Lim LY, Tsai HY, Kuo WC, Tsai PH, Yang SH, Hong SI, Chen SJ, Hwang DK, Yang YP, Chiou SH. Inhibition of oxidative stress-induced epithelial-mesenchymal transition in retinal pigment epithelial cells of age-related macular degeneration model by suppressing ERK activation. J Adv Res. 2023 Jun 15;60:141–57. doi:10.1016/j.jare.2023.06.004 PubMed PMID: 37328058; PubMed Central PMCID: PMC11156608.

40. Chen T, You Y, Jiang H, Wang ZZ. Epithelial-mesenchymal transition (EMT): A biological process in the development, stem cell differentiation and tumorigenesis. J Cell Physiol. 2017 Dec;232(12):3261–72. doi:10.1002/jcp.25797 PubMed PMID: 28079253; PubMed Central PMCID: PMC5507753.

41. Shu DY, Butcher E, Saint-Geniez M. EMT and EndMT: Emerging Roles in Age-Related Macular Degeneration. Int J Mol Sci. 2020 Jun 16;21(12):4271. doi:10.3390/ijms21124271 PubMed PMID: 32560057; PubMed Central PMCID: PMC7349630.

42. Hill C, Jones MG, Davies DE, Wang Y. Epithelial-mesenchymal transition contributes to pulmonary fibrosis via aberrant epithelial/fibroblastic cross-talk. J Lung Health Dis. 2019 Apr 2;3(2):31–5. PubMed PMID: 31032489; PubMed Central PMCID: PMC6485666.

43. Yeung KT, Yang J. Epithelial–mesenchymal transition in tumor metastasis. Mol Oncol. 2017 Jan;11(1):28–39. doi:10.1002/1878-0261.12017 PubMed PMID: 28085222; PubMed Central PMCID: PMC5242415.

44. Huang Y, Hong W, Wei X. The molecular mechanisms and therapeutic strategies of EMT in tumor progression and metastasis. J Hematol Oncol. 2022 Sep 8;15:129. doi:10.1186/s13045-022-01347-8 PubMed PMID: 36076302; PubMed Central PMCID: PMC9461252.

45. Dave N, Guaita-Esteruelas S, Gutarra S, Frias À, Beltran M, Peiró S, García de Herreros A. Functional Cooperation between Snail1 and Twist in the Regulation of ZEB1 Expression during Epithelial to Mesenchymal Transition. J Biol Chem. 2011 Apr 8;286(14):12024–32. doi:10.1074/jbc.M110.168625 PubMed PMID: 21317430; PubMed Central PMCID: PMC3069405.

46. Niknami Z, Muhammadnejad A, Ebrahimi A, Harsani Z, Shirkoohi R. Significance of E-cadherin and Vimentin as epithelial-mesenchymal transition markers in colorectal carcinoma prognosis. EXCLI J. 2020 Jun 26;19:917–26. doi:10.17179/excli2020-1946 PubMed PMID: 32665775; PubMed Central PMCID: PMC7355153.

47. Datta S, Cano M, Satyanarayana G, Liu T, Wang L, Wang J, Cheng J, Itoh K, Sharma A, Bhutto I, Kannan R, Qian J, Sinha D, Handa JT. Mitophagy initiates retrograde mitochondrial-nuclear signaling to guide retinal pigment cell heterogeneity. Autophagy. 2023 Mar;19(3):966–83. doi:10.1080/15548627.2022.2109286 PubMed PMID: 35921555; PubMed Central PMCID: PMC9980637.

48. Singh K, Jin Y, Hu MW, Palazzo I, Cano M, Hoang T, Bhutto I, Wang S, Sinha D, Blackshaw S, Qian J, Handa JT. Cigarette smoke and biological age induce degenerative heterogeneity in retinal pigment epithelium [Internet]. bioRxiv; 2025 [cited 2025 Sep 21]. p. 2025.01.27.635096. Available from: [https://www.biorxiv.org/content/10.1101/2025.01.27.635096v1](https://www.biorxiv.org/content/10.1101/2025.01.27.635096v1)doi:10.1101/2025.01.27.635096

49. Fisher CR, Ferrington DA. Perspective on AMD Pathobiology: A Bioenergetic Crisis in the RPE. Invest Ophthalmol Vis Sci. 2018 Jun;59(4):AMD41–7. doi:10.1167/iovs.18-24289 PubMed PMID: 30025108; PubMed Central PMCID: PMC5989860.

50. Yu AL, Birke K, Burger J, Welge-Lussen U. Biological Effects of Cigarette Smoke in Cultured Human Retinal Pigment Epithelial Cells. PLoS One. 2012 Nov 14;7(11):e48501. doi:10.1371/journal.pone.0048501 PubMed PMID: 23155386; PubMed Central PMCID: PMC3498276.

51. Wang L, Kaya KD, Kim S, Brooks MJ, Wang J, Xin Y, Qian J, Swaroop A, Handa JT. Retinal pigment epithelium transcriptome analysis in chronic smoking reveals a suppressed innate immune response and activation of differentiation pathways. Free Radic Biol Med. 2020 Aug 20;156:176–89. doi:10.1016/j.freeradbiomed.2020.06.004 PubMed PMID: 32634473; PubMed Central PMCID: PMC7434665.

52. Wang L, Kondo N, Cano M, Ebrahimi K, Yoshida T, Barnett BP, Biswal S, Handa JT. Nrf2 Signaling Modulates Cigarette Smoke Induced Complement Activation in Retinal Pigmented Epithelial Cells. Free Radic Biol Med. 2014 May;70:155–66. doi:10.1016/j.freeradbiomed.2014.01.015 PubMed PMID: 24440594; PubMed Central PMCID: PMC4006310.

53. Burke JM. Epithelial phenotype and the RPE: is the answer blowing in the Wnt? Prog Retin Eye Res. 2008 Nov;27(6):579–95. doi:10.1016/j.preteyeres.2008.08.002 PubMed PMID: 18775790; PubMed Central PMCID: PMC2584165.

54. Burke JM, Skumatz CM, Irving PE, McKay BS. Phenotypic heterogeneity of retinal pigment epithelial cells in vitro and in situ. Exp Eye Res. 1996 Jan;62(1):63–73. doi:10.1006/exer.1996.0008 PubMed PMID: 8674514.

55. Ghosh S, Shang P, Terasaki H, Stepicheva N, Hose S, Yazdankhah M, Weiss J, Sakamoto T, Bhutto IA, Xia S, Zigler JS Jr, Kannan R, Qian J, Handa JT, Sinha D. A Role for βA3/A1-Crystallin in Type 2 EMT of RPE Cells Occurring in Dry Age-Related Macular Degeneration. Invest Ophthalmol Vis Sci. 2018 Mar 20;59(4):AMD104–13. doi:10.1167/iovs.18-24132 PubMed PMID: 30098172; PubMed Central PMCID: PMC6058694.

56. Zanzottera EC, Messinger JD, Ach T, Smith RT, Curcio CA. Subducted and melanotic cells in advanced age-related macular degeneration are derived from retinal pigment epithelium. Invest Ophthalmol Vis Sci. 2015 May;56(5):3269–78. doi:10.1167/iovs.15-16432 PubMed PMID: 26024109; PubMed Central PMCID: PMC4453434.

57. Auciello FR, Ross FA, Ikematsu N, Hardie DG. Oxidative stress activates AMPK in cultured cells primarily by increasing cellular AMP and/or ADP. FEBS Lett. 2014 Sep 17;588(18):3361–6. doi:10.1016/j.febslet.2014.07.025 PubMed PMID: 25084564; PubMed Central PMCID: PMC4158911.

58. Kaarniranta K, Blasiak J, Liton P, Boulton M, Klionsky DJ, Sinha D. Autophagy in age-related macular degeneration. Autophagy. 2023 Feb;19(2):388–400. doi:10.1080/15548627.2022.2069437 PubMed PMID: 35468037; PubMed Central PMCID: PMC9851256.

59. Jiménez-Loygorri JI, Viedma-Poyatos Á, Gómez-Sintes R, Boya P. Urolithin A promotes p62-dependent lysophagy to prevent acute retinal neurodegeneration. Mol Neurodegener. 2024 Jun 18;19:49. doi:10.1186/s13024-024-00739-3 PubMed PMID: 38890703; PubMed Central PMCID: PMC11186080.

