## Supplementary Figure Legends for "Effect of Varying Cigarette Smoke Concentrations and Durations on the Retinal Pigment Epithelium"

### SUPPLEMENTAL MATERIAL

**Supplemental Table 1.** TaqMan (Thermo Fisher Scientific) primer probes used for RT-qPCR analysis.

**Supplemental Table 2.** Primary antibodies used for western blot analysis.

**Supplemental Figure 1.** CSC induces dose-dependent increase in mitochondrial superoxide levels at increasing concentrations in A. ARPE-19 cells treated for 4 hours, B. ARPE-19 cells treated for 24 hours, C. MRPE cells treated for 4 hours, and D. MRPE cells treated for 24 hours. MitoSOX Red fluorescently labels mitochondrial superoxide red; Hoechst staining fluorescently labels cell nuclei blue. Quantification plots are shown in Figure 1.

**Supplemental Figure 2.** CSC induces morphological changes at increasing concentrations of CSC at 4 and 24 hours. A. ARPE-19 cells treated 4 hours, B.ARPE-19 cells treated for 24 hours, C. MRPE cells treated for 4 hours, and D. MRPE cells treated for 24 hours. RPE morphology plots are shown in Supplemental Figure 3.

**Supplemental Fig 3.** RPE Morphometry in different cell lines and treatment groups. Aspect ratio is represented in mean ± SD, Student’s t-test; *p < 0.05, **p < 0.01 ***p < 0.001, ****p < 0.0001.

**Supplemental Fig 4.** CSC induces concentration-dependent increase in cell death at increasing concentrations in A. ARPE-19 cells treated for 4 hours, B. ARPE-19 cells treated for 24 hours, C. MRPE cells treated for 4 hours, and D. MRPE cells treated for 24 hours. Red-fluorescent ethidium homodimer selectively detects dead cells; Hoechst staining fluorescently labels cell nuclei blue. Quantification plots are shown in Figure 5.

**Supplementary Table 3.** Reference for gene and protein nomenclature.
