## Supplementary Table 1 for "Effect of Varying Cigarette Smoke Concentrations and Durations on the Retinal Pigment Epithelium"

| **Gene symbol** | **Species** | **Source** | **Assay Type** | **Assay ID / Catalog no.** |
| --- | --- | --- | --- | --- |
| *Gapdh* | Mouse | ThermoFisher Scientific | TaqMan | Mm99999915 |
| *Map1lc3b* | Mouse | ThermoFisher Scientific | TaqMan | Mm00782868 |
| *Zeb1* | Mouse | ThermoFisher Scientific | TaqMan | Mm00495564 |
| *Snai1* | Mouse | ThermoFisher Scientific | TaqMan | Mm00441533 |
| *Sqstm1* | Mouse | ThermoFisher Scientific | TaqMan | Mm00448091 |
| *Nfe2l2* | Mouse | ThermoFisher Scientific | TaqMan | Mm00477784 |
| *GAPDH* | Human | ThermoFisher Scientific | TaqMan | Hs99999905 |
| *NFE2L2* | Human | ThermoFisher Scientific | TaqMan | Hs00975960 |
| *ZEB1* | Human | ThermoFisher Scientific | TaqMan | Hs01566408 |
| *SNAI1* | Human | ThermoFisher Scientific | TaqMan | Hs00195591 |
| *SQSTM1* | Human | ThermoFisher Scientific | TaqMan | Hs00177654 |
| *MAP1LC3B* | Human | ThermoFisher Scientific | TaqMan | Hs00797944 |

**Supplemental Table 1.** TaqMan (Thermo Fisher Scientific) primer probes used for RT-qPCR analysis.
