## Supplementary Table 2 for "Effect of Varying Cigarette Smoke Concentrations and Durations on the Retinal Pigment Epithelium"

| **Antibody** | **Company** | **Dilution** |
| --- | --- | --- |
| LC3B rabbit mAb | Millipore | 1:1000 |
| P62 mouse mAb (M162-3) | MBL | 1:1000 |
| P62 rabbit pAb (PM045) | MBL | 1:1000 |
| E-cadherin rabbit mAb (24E10) | Cell Signaling Technologies | 1:1000 |
| Vimentin mouse mAb (sc-373717) | Santa Cruz | 1:1000 |
| Beta-actin mouse mAb-HRP (sc-47778) | Santa Cruz | 1:1000 |
| NRF2 rabbit mAb (D1Z9C) | Cell Signaling Technologies | 1:1000 |
| AMPK rabbit (#5831S) | Cell Signaling Technologies | 1:1000 |
| pAMPK rabbit (#2535S) | Cell Signaling Technologies | 1:1000 |
| Goat anti-rabbit IgG (HRP) (ab6721) | Abcam | 1:2000 |
| Rabbit anti-mouse IgG (HRP) (ab6728) | Abcam | 1:2000 |

**Supplemental Table 2.** Primary antibodies used for western blot analysis.
