## Supplementary Table 3 for "Effect of Varying Cigarette Smoke Concentrations and Durations on the Retinal Pigment Epithelium"

| **Target / Marker** | **Human Gene Name** | **Mouse Gene Name** | **Human/Mouse Protein Name** |
| --- | --- | --- | --- |
| LC3 (LC3B) | *MAP1LC3B* | *Map1lc3b* | LC3B |
| p62 | *SQSTM1* | *Sqstm1* | P62 |
| Nrf2 | *NFE2L2* | *Nfe2l2* | NRF2 |
| E-cadherin | *CDH1* | *Cdh1* | E-CADHERIN |
| Vimentin | *VIM* | *Vim* | VIMENTIN |
| AMPK (α1) | *PRKAA1* | *Prkaa1* | AMPK |
| p-AMPK | *N/A* | *N/A* | P-AMPK |
| ZEB1 | *ZEB1* | *Zeb1* | ZEB1 |
| SNAI1(Snail) | *SNAI1* | *Snai1* | SNAI1 |

**Supplemental Table 3.** Reference for gene and protein nomenclature
