## Supplementary Figures for "Effect of Varying Cigarette Smoke Concentrations and Durations on the Retinal Pigment Epithelium"

**Supplemental Table 3.** Reference for gene and protein nomenclature

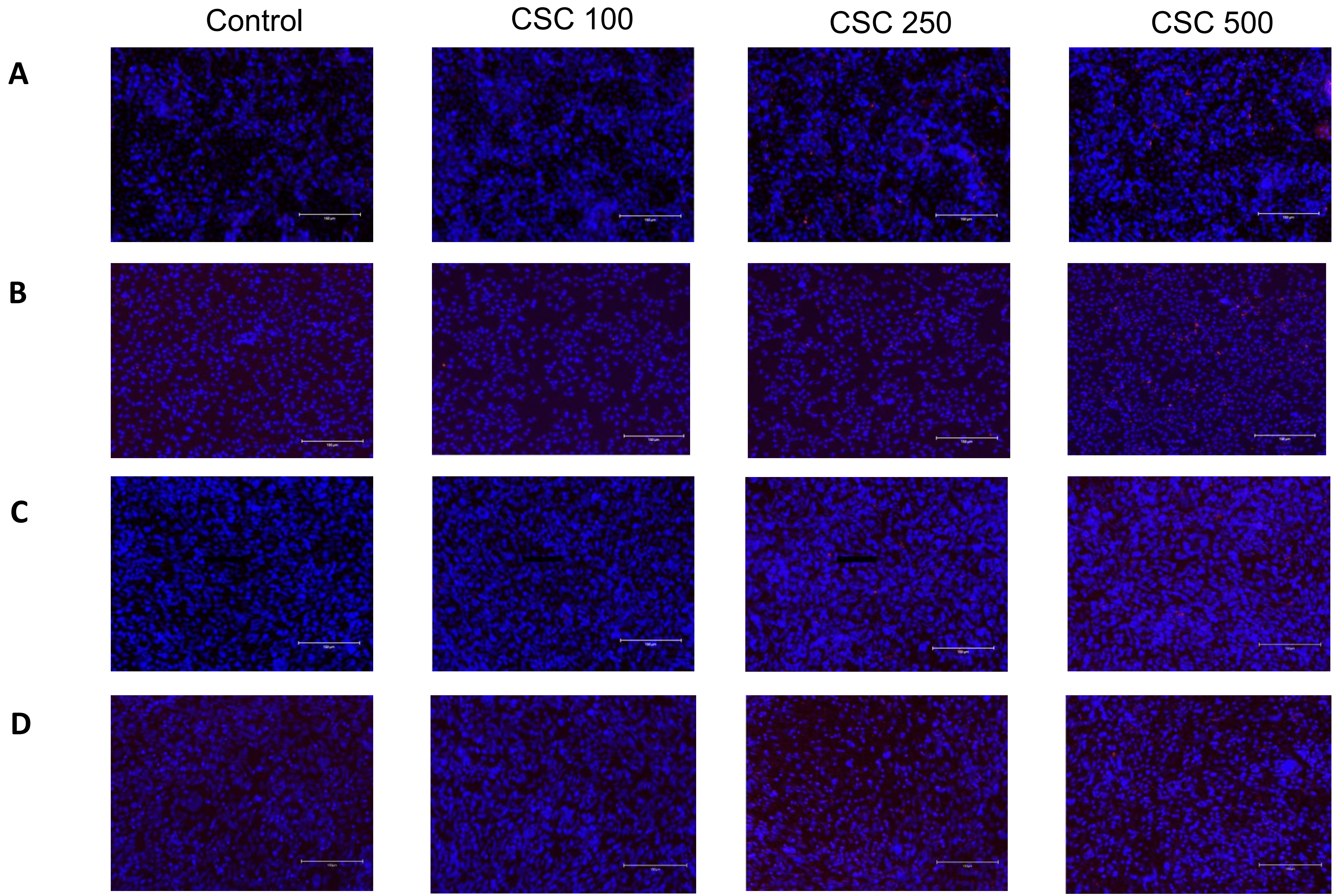


**Supplemental Figure 1**. CSC induces dose-dependent increase in mitochondrial superoxide levels at increasing concentrations in A. ARPE-19 cells treated for 4 hours, B. ARPE-19 cells treated for 24 hours, C. MRPE cells treated for 4 hours, and D. MRPE cells treated for 24 hours. MitoSOX Red fluorescently labels mitochondrial superoxide red; Hoechst staining fluorescently labels cell nuclei blue. Quantification plots are shown in Figure 1.

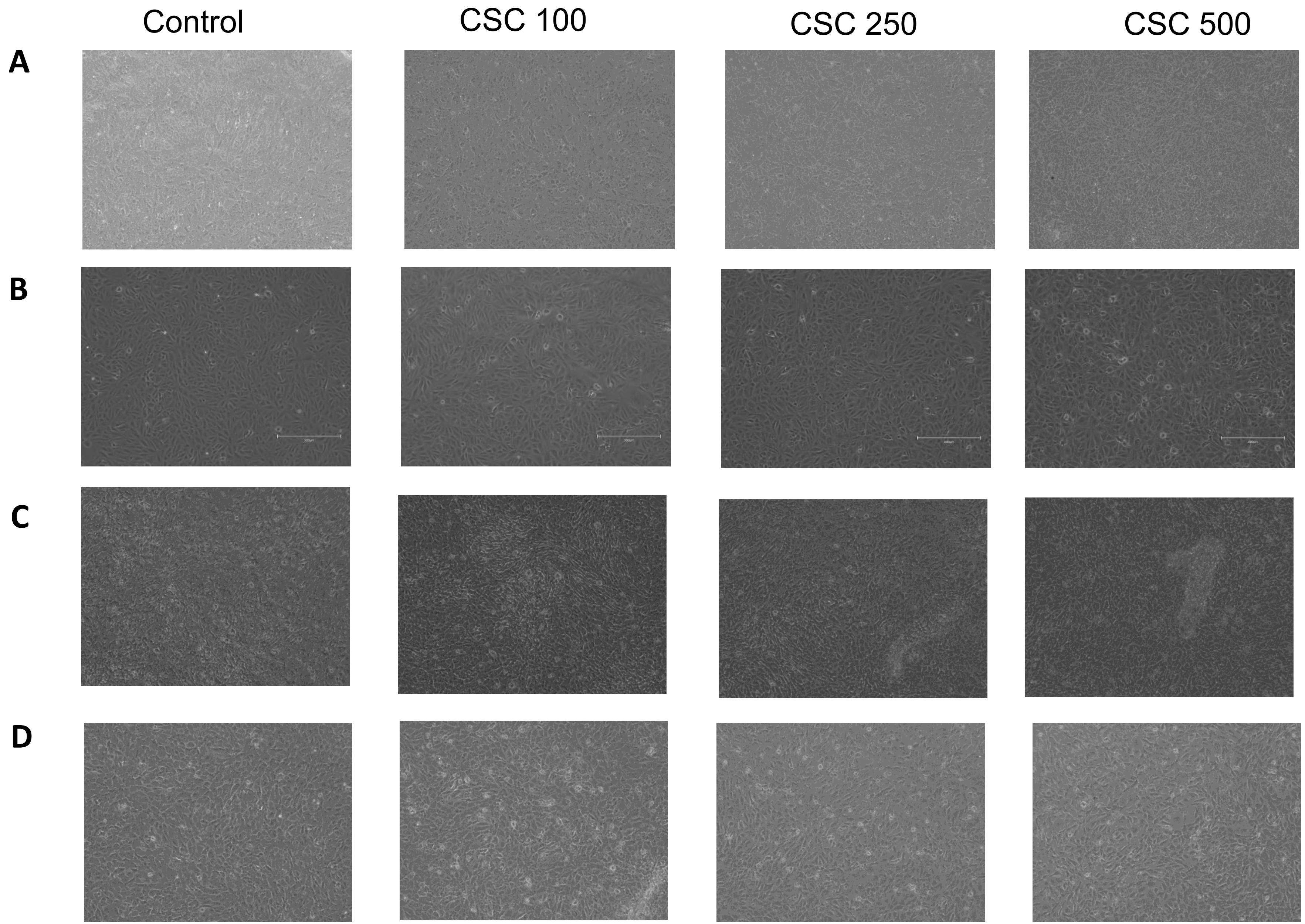


**Supplemental Figure 2**. CSE induces morphological changes at increasing doses of CSC at 4 and 24 hours. A. ARPE-19 cells treated with CSE for 4 hours, B. ARPE-19 cells treated with CSE for 24 s, C. MRPE cells treated with CSE for 4 hours, and D. MRPE cells treated with CSE for 24 hours. RPE morphology plots are included in supplemental Figure 4.

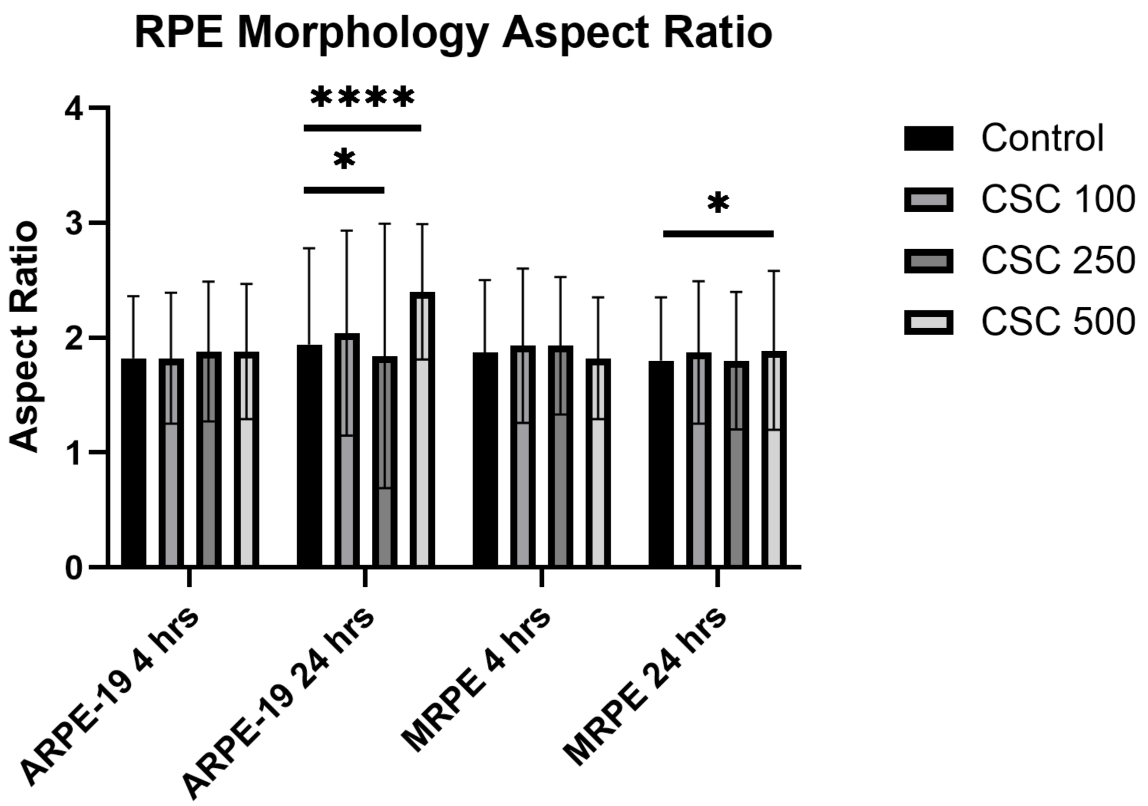


**Supplemental Figure 3**. RPE Morphometry in different cell lines and treatment groups. Aspect ratio is represented in mean ± SD, Student's t-test; *P < 0.05, **P < 0.01 ***P < 0.001.

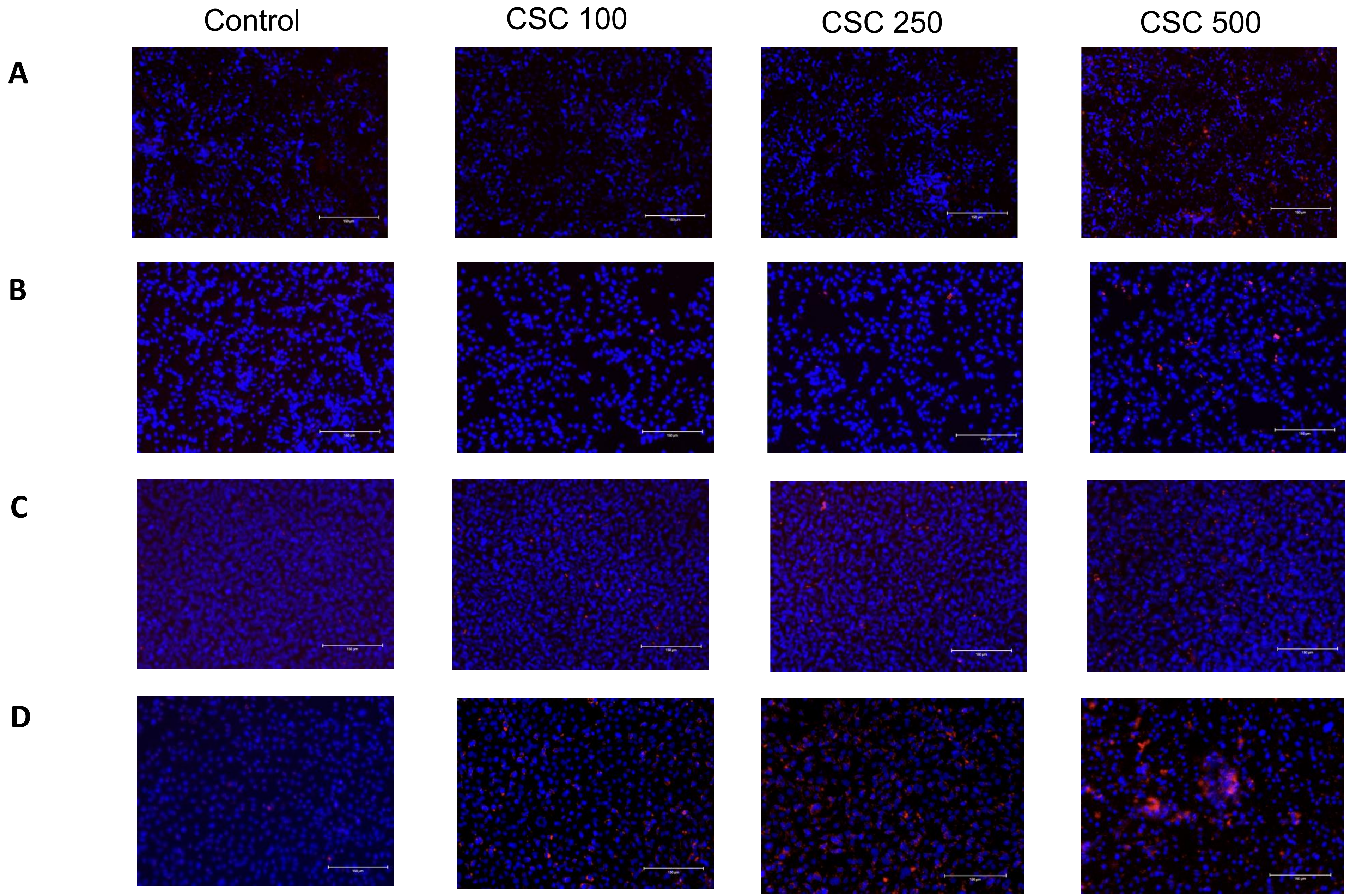


**Supplemental Figure 4**. CSE induces dose-dependent increase in cell death at increasing doses in A. ARPE-19 cells treated with CSE for 4 hours, B. ARPE-19 cells treated with CSE for 24 hours, C. MRPE cells treated with CSE for 4 hours, and D. MRPE cells treated with CSE for 24 hours. Red-fluorescent ethidium homodimer selectively detects dead cells; Hoescht staining fluorescently labels cell nuclei blue.  Quantification plots are included in Figure 5.
